# Mechanisms of improved specificity of engineered Cas9s revealed by single molecule analysis

**DOI:** 10.1101/192724

**Authors:** Digvijay Singh, Yanbo Wang, John Mallon, Olivia Yang, Jingyi Fei, Anustup Poddar, Damon Ceylan, Scott Bailey, Taekjip Ha

## Abstract

In microbes, CRISPR-Cas systems provide adaptive immunity against invading genetic elements. Cas9 in complex with a guide-RNA targets complementary DNA for cleavage and has been repurposed for wide-ranging biological applications. New Cas9s have been engineered (eCas9 and Cas9-HF1) to improve specificity, but how they help reduce off-target cleavage is not known. Here, we developed single molecule DNA unwinding assay to show that sequence mismatches affect cleavage reactions through rebalancing the internal unwinding/rewinding equilibrium. Increasing PAM-distal mismatches facilitate rewinding, and the associated cleavage impairment shows that cleavage proceeds from the unwound state. Engineered Cas9s depopulate the unwound state more readily upon mismatch detection. Intrinsic cleavage rate is much lower for engineered Cas9s, preventing cleavage from transiently unwound off-targets. DNA interrogation experiments showed that engineered Cas9s require about one additional base pair match for stable binding, freeing them from sites that would otherwise sequester them. Therefore, engineered Cas9s achieve their improved specificity (1) by inhibiting stable DNA binding to partially matching sequences, (2) by making DNA unwinding more sensitive to mismatches, and (3) by slowing down intrinsic cleavage reaction.

---

In bacteria and archaea, CRISPR (clustered regularly interspaced short palindromic repeats)–Cas systems impart adaptive defense against phages and plasmid^1^. In type II CRISPR-Cas systems, the Cas9 endonuclease in complex with two RNAs, a CRISPR guide-RNA (crRNA) and a trans-activating RNA (tracrRNA), targets complementary 20 base pair (bp) sequences (protospacers) in foreign DNA for double-stranded cleavage, with a requirement that they be followed by a motif called PAM (protospacer adjacent motif, 5_’_-NGG-3_’_ for *S. pyogenes* Cas9)^2-4^. Programmable DNA binding and cleavage by Cas9 has revolutionized life sciences, where Cas9 cleavage is used for genome editing and Cas9 binding is used for tagging genomic sites with markers or effectors for wide-ranging applications^5,6^. Minimizing off-target effects of both binding and cleavage^7,8^ remains an active area of study. *In vitro* and *in vivo* investigations have shown that Cas9-RNA specificity is most affected by PAM plus an 8-10 PAM-proximal seed region and concentrations of both Cas9 and RNA^7,9^. Rationally designed and engineered *S. Pyogenes* Cas9s (EngCas9s), namely enhanced Cas9 (eCas9; K848A/K1003A/R1060A mutations) and high fidelity Cas9 (Cas9-HF1; N497A/R661A/Q695A/Q926A mutations), led to remarkable specificity improvements in unbiased genome wide CRISPRCas9 specificity measurements^10,11^.

The stability of Cas9-RNA-DNA complex is in part a function of the specific RNA-DNA base-pairing and sequence-independent interactions between Cas9 residues and DNA, the latter being a potential source of promiscuity. Cas9-RNA uses binding energy to initially melt DNA near PAM, and DNA unwinding can continue downstream in the presence of sequence complementary with guide-RNA. The motivation behind the mutations in EngCas9s was to destabilize the Cas9-RNA-DNA complex by diminishing the favorable sequence-independent interactions between Cas9 residues and the DNA backbone (**Fig. 1b**). How these mutations affect and modulate DNA interrogation, unwinding and cleavage remains unknown. Here, we report a comparative analysis of DNA interrogation and rejection, DNA unwinding and rewinding dynamics, and cleavage activation among different Cas9s using single molecule imaging methods that can detect multiple conformations and transient intermediates^12^ and have been used previously to study CRISPR systems *in vitro*^9,13-21^.

**Figure 1.**
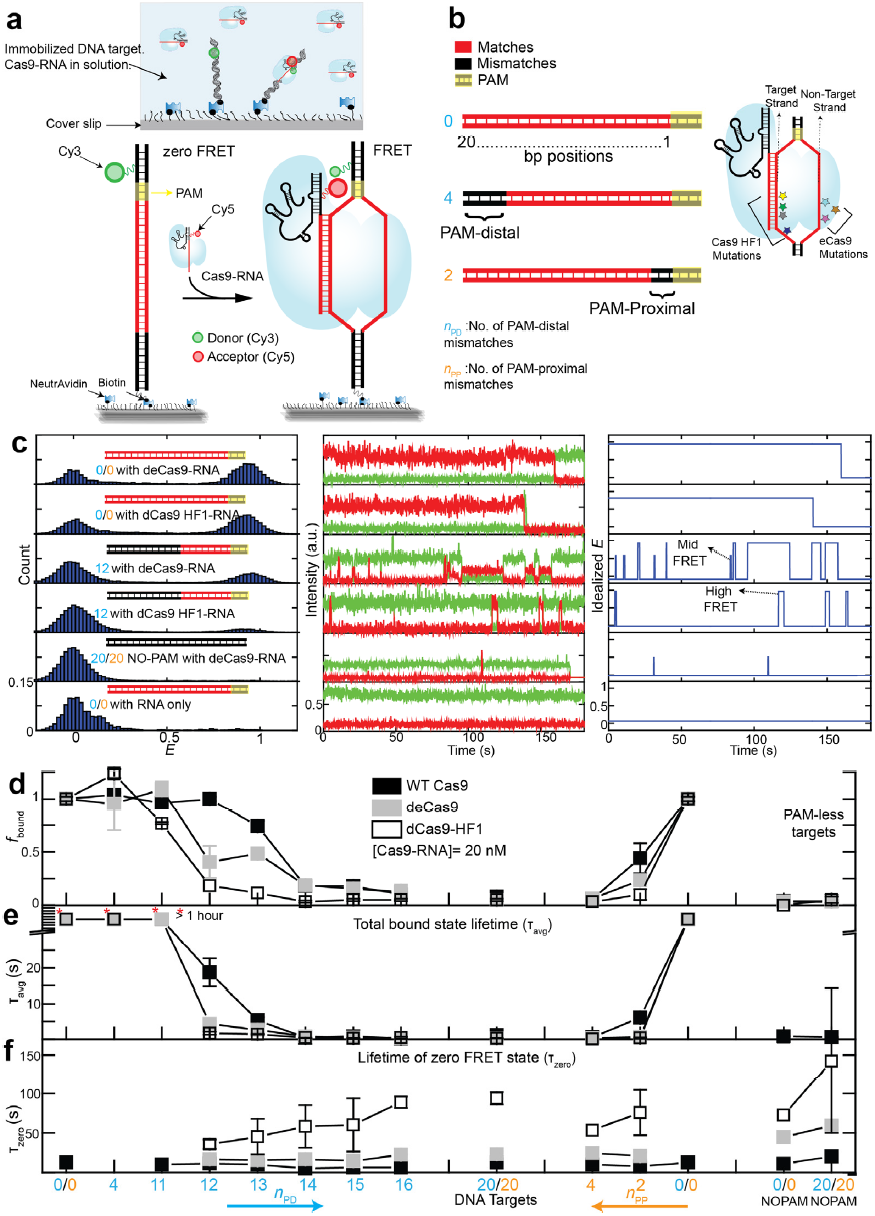
smFRET assay to study DNA interrogation by engineered Cas9-RNA. **(a)** Schematic of smFRET assay. Cas9 in complex with an acceptor labeled guide-RNA binds a donor-labeled cognate DNA target. (**b)** DNA targets with mismatches in the protospacer region against the guide-RNA. The number of mismatches PAM-distal (*n*_PD_) and PAM-proximal (*n*_PP_) are shown in cyan and orange, respectively. Also shown are locations of EngCas9 mutations in dCas9-RNA-DNA complex (PDB ID: 4UN3). **(c)** *E* histograms (left) at 20 nM EngCas9-RNA or RNA only. Representative single molecule intensity time traces of donor (green) and acceptor (red) are shown (middle), along with *E* values idealized (right) by hidden Markov modeling. **(d)** Normalized fraction (*f*_bound_) of DNA molecules bound with Cas9-RNA at 20 nM Cas9-RNA. Fractions were normalized relative to the bound fraction of cognate DNA target. **(e)** *τ*_avg_, obtained from dwell times of *E* >0.2 states. **(f)** Unbound state lifetimes at [Cas9-RNA] = 20 nM. The mean of *τ*_avg_ over various DNA targets was used to calculate *k*_on_. Number of PAM-distal (*n*_PD_) and PAM-proximal mismatches (*n*_PP_) are shown in cyan and orange respectively. Error bars represent s.d. Data for WT Cas9-RNA is taken from a previous study^9^.

## Results

### Real-time single molecule FRET assay for DNA interrogation

We employed a single-molecule fluorescence resonance energy transfer (smFRET) assay^22^ to investigate DNA interrogation by EngCas9-RNA. DNA targets (donor-labeled, 82 bp long) were immobilized on polyethylene glycol (PEG)-passivated flow chambers and EngCas9 pre-complexed with acceptor-labeled guide-RNA (EngCas9-RNA) was added to observe their interactions in real time. Locations of donor (Cy3) and acceptor (Cy5) were chosen such that specific interaction between DNA target and EngCas9-RNA would lead to FRET (**Fig. 1a** and **Supplementary Fig. 1**). Labeling at these locations do not affect Cas9 cleavage activity^9^. Unless mentioned otherwise, we used catalytically dead versions of all Cas9s (denoted with the prefix d) in order to focus on the properties prior to cleavage. Wild-type (WT) Cas9 and dCas9 showed similar behavior in DNA interrogation experiments^9^.

We examined different DNA targets containing mismatches relative to the guide-RNA. As the naming convention, we used *n*_PD_ (the number of PAM-distal mismatches) and *n*_PP_ (the number of PAM-proximal mismatches) (**Fig. 1b**). The cognate DNA target gave two distinct populations centered at FRET efficiency (*E)* of 0.9 and 0 in the presence of 20 nM EngCas9-RNA. The *E*=0.9 population was negligible if only the labeled RNA was added without Cas9 or if a fully mismatched DNA without PAM was used. Therefore, we assigned the *E*=0.9 species to a sequence-specific EngCas9-RNA-DNA complex (**Fig. 1c**). The *E*=0 population is a combination of unbound states and bound states with inactive or missing acceptor. Cas9-RNA titration gave the apparent dissociation constant (*K*_*d*_) of 0.50 nM (dCas9), 0.5 nM (deCas9) and 2.7 nM (dCas9-HF1) (**Supplementary Fig. 2**).

To quantify the impact of mismatches, we determined the apparent Cas9-RNA bound fraction *f*_bound_, defined as the normalized fraction of DNA molecules with *E* > 0.75 (20 nM Cas9-RNA) (**Fig. 1d** and **Supplementary Fig. 3**). *f*_bound_ vs. mismatches for EngCas9s was similar to that observed previously with WT Cas9^9^; (i) *f*_bound_ remained unchanged when *n*_PD_ increased from 0 to 10 or 11, but precipitously decreased beyond, (ii) even *n*_PP_ of 2 or 4 caused a >50 % or >95% drop in *f*_bound_, respectively, (iii) binding is ultra-stable with >8-9 PAM-proximal matches as *f*_bound_ remained high even 1 hour after washing away free Cas9-RNA (**Supplementary Fig. 4**).

If there are enough mismatches to preclude stable binding, smFRET time-traces showed repetitive transitions between *E*=0 and *E*=0.45 states in addition to transitions between *E*=0 and *E*=0.9, suggesting that there are multiple bound states distinguishable based on *E* ^9^ (**Fig. 1c**). We used a hidden Markov modeling analysis^23^ to determine *τ*_avg_ as a fraction-weighted average of the high (*E*=0.9) and mid (*E*=0.45) FRET state lifetimes. *τ*_avg_ was over 1 hour for DNA targets with at least *m* PAM-proximal matches (*m*=9 for WT Cas9 and deCas9, 10 for dCas9-HF1) but decreased to 0.5 – 15 s with fewer than *m* PAM-proximal matches or any PAM-proximal mismatches (**Fig. 1e**). Dwell times of the unbound state (*E* <0.2) were only weakly dependent on sequence (**Fig. 1f**) (**Supplementary Fig. 5-6**)

We list below qualitative features common between EngCas9s and WT Cas9 as well as quantitative differences:

(1) All Cas9s interrogate and bind DNA in two distinct modes. Sequence-independent sampling of DNA target in search of PAM results in transient mid FRET events. The high FRET state results upon PAM detection and RNA-DNA heteroduplex formation, and its lifetime increased with increasing base-pairing between guide-RNA and DNA target. (2) The binding frequency is independent of sequence. The bimolecular association rate constant *k*_on_ decreased for EngCas9s (2.6×10^6^ M^-1^ s^-1^ for deCas9 and 0.9×10^6^ M-^1^ s^-1^ for dCas9-HF1, compared to 5.4×10^6^ M^-1^ s^-1^ for WT Cas9), possibly as a result of alteration in electrostatic and polar interaction upon removal of positively charged and polar residues (**Fig. 1f**). (3) PAM-proximal mismatches are much more deleterious for stable binding compared to PAM-distal mismatches, and mismatches in the middle of the protospacer renders PAM-distal matches inconsequential (**Supplementary Fig. 3**) ^9^, suggesting that all Cas9s extend RNA-DNA heteroduplex unidirectionally from PAM-proximal to PAM-distal end. (4) PAM-proximal mismatches are more deleterious for EngCas9 binding than WT Cas9, and (5) deCas9 and WT Cas9^9,24^ require 9 PAM-proximal matches for ultra-stable binding whereas dCas9-HF1 requires 10 (**Fig. 1e**, **Supplementary Fig. 4**). Overall, EngCas9 are similar to WT Cas9 in their sequence-dependent binding properties except for the higher ability to reject both PAM-proximal and PAM-distal mismatches which may help reduce Cas9-RNA sequestration by partially matching targets.

### Cas9-RNA induced DNA unwinding

Genome wide characterization of EngCas9 cleavage specificity showed marked improvements over WT Cas9^10,11^. Yet, our binding study shows that the EngCas9s still stably bind to DNA with *n*_PD_ as large as 10. We reasoned that EngCas9s may differ from WT Cas9 in their mismatch dependence of DNA unwinding which, if promoted by annealing to guide-RNA, may be a determinant of cleavage action^25^. Therefore, we probed the internal unwinding of PAM-distal region of the protospacer by labeling the target and non-target strands with a donor and an acceptor, respectively, with 9 bp spacing (**Supplementary Fig. 7**). Labeling at these locations did not perturb cleavage (**Supplementary Fig. 8**). The DNA by itself showed high FRET (*E~*0.75) (**Fig. 2a**), and upon addition of 100 nM WT dCas9-RNA to cognate DNA, we observed a shift to a stable low FRET state (*E~*0.30), likely because of DNA unwinding (**Fig. 2b**). A similar FRET change was observed when the locations of the donor and acceptor were swapped (**Supplementary Fig. 7b**). A DNA with *n*_PD_ = 1 showed a similarly stable unwinding but with a slightly higher *E* of ~0.35 and occasional short-lived transitions to the high FRET state, likely due to one bp fewer unwinding and increased frequency to rewind, respectively. The trend continued upon increasing *n*_PD_ to 2 and 3; *E* value for the unwound state increased, and the relative population of the rewound state increased. With *n*_PD_ = 4, the unwound state is rarely populated. deCas9 and dCas9-HF1 showed a similar behavior but their unwound state population and lifetime were smaller and decreased more quickly with increasing *n*_PD_ (**Fig. 2b**). Therefore, EngCas9s are more sensitive to PAM-distal mismatches in their ability to unwind DNA. The rewound state must still have at least 8 PAM-proximal bp unwound because we did not see stable binding with fewer PAM-proximal matches (**Supplementary Fig. 4**), and can have up to 16 PAM-proximal bp unwound.

**Figure 2.**
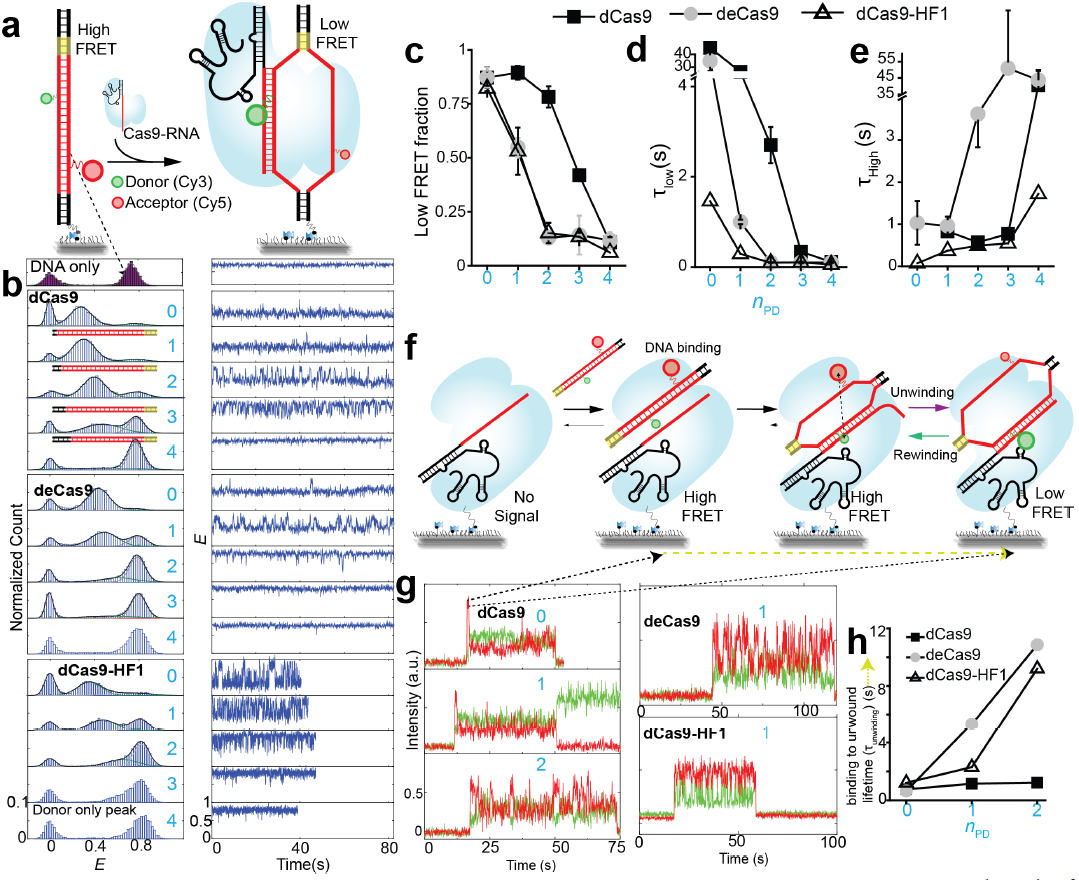
Internal DNA unwinding/rewinding dynamics modulated by mismatches and Cas9 mutations. **(a)** Schematic of smFRET assay for Cas9-RNA induced unwinding of surface-tethered DNA. **(b)** *E* histograms for *n*_PD_ = 0, 1, 2, 3 and 4 (cyan numbering) and representative time traces. **(c-e)** Relative population of the unwound state, average lifetime of the unwound state and average lifetime of the rewound state vs. *n*_PD_. **(f)** Schematic of smFRET assay for DNA unwinding by surface-tethered Cas9-RNA. **(g)** Representative single molecule fluorescence intensity time traces of donor (green) and acceptor (red) show abrupt signal increase upon DNA binding followed by FRET changes. **(h)** Time taken to go from initial binding to its first unwound state configuration (*τ*_unwinding_) as denoted by a dark yellow line (f).

We observed two distinct *E* populations even at a higher time resolution (35 ms) (**Supplementary Fig. 9**), indicating that Cas9-RNA induces primarily two states, unwound and rewound, without spending appreciable time in between. We used the Hidden Markov modeling analysis to segment single molecule time traces into two states (**Supplementary Fig. 10**). All Cas9s showed a reduction in the relative population and lifetime of the unwound state with increasing *n*_PD_ (**Fig. 2c-e**). Combined with the gradual increase of *E* values for the unwound state (**Fig. 2b**), we can conclude that PAM-distal mismatches reduce the time spent in the unwound state in addition to reducing the maximal extent of unwinding. For a given nPD, EngCas9s showed lower occupancy and shorter lifetime of the unwound state (**Fig. 2c-e**), suggesting that the mutations indeed destabilize the maximally unwound states. We observed a similar unwinding behavior from catalytically active Cas9 but *E* distribution was broader and state transitions were less frequent (**Supplementary Fig. 11**), suggesting a change in unwinding dynamics after cleavage, possibly due to disordered non-target strand^25,26^.

In order to capture the initial DNA unwinding event, we added labeled DNA to surface-immobilized Cas9-RNA molecules (**Fig. 2f**) (**Supplementary Fig. 12-13**). DNA binding is detected as a sudden appearance of fluorescence signal in a high *E* state, followed by a single step change to a low *E* (unwound state) (**Fig. 2g**). Subsequently, we observed unwinding/rewinding dynamics similar to what we observed in the steady state (**Fig. 2g**) (**Supplementary Fig. 14-15**). The average dwell time of the initial high *E* state which we attribute to the time it takes to unwind DNA for the first time (*τ*_unwinding_) remained constant (~1 s) for dCas9 when *n*_PD_ changed from 0 to 2. In contrast, *τ*_unwinding_ increased ~9 folds for EngCas9s when *n*_PD_ changed from 0 to 2 (**Fig. 2h**), suggesting that EngCas9s take longer to unwind in the presence of PAM-distal matches.

### DNA cleavage vs. mismatches

Next, we probed how mismatches influence target cleavage through modulating DNA unwinding/rewinding dynamics. Cleavage was more specific for EngCas9s because they cleaved only up to nPD = 3 compared to *n*_PD_ = 4 for WT Cas9 (**Fig. 3a & Supplementary Fig. 8**). EngCas9s did not release cleavage products under physiological conditions (**Supplementary Fig. 4**), as was the case for WT Cas9^9^. In terms of the cleavage reaction, EngCas9s were much slower than WT Cas9 for cognate or PAM-distal mismatched DNA (**Fig. 3b-c**) but were faster for PAM-proximal mismatched DNA (**Fig. 3a**). Overall, we observed a clear correlation between the cleavage time scale, *τ*_cleavage_, and DNA unwinding signatures; *τ*_cleavage_ decreased with increasing population (**Fig. 3d**) and lifetime (**Fig. 3e**) of the unwound state. Therefore, cleavage must proceed from the unwound state.

**Figure 3.**
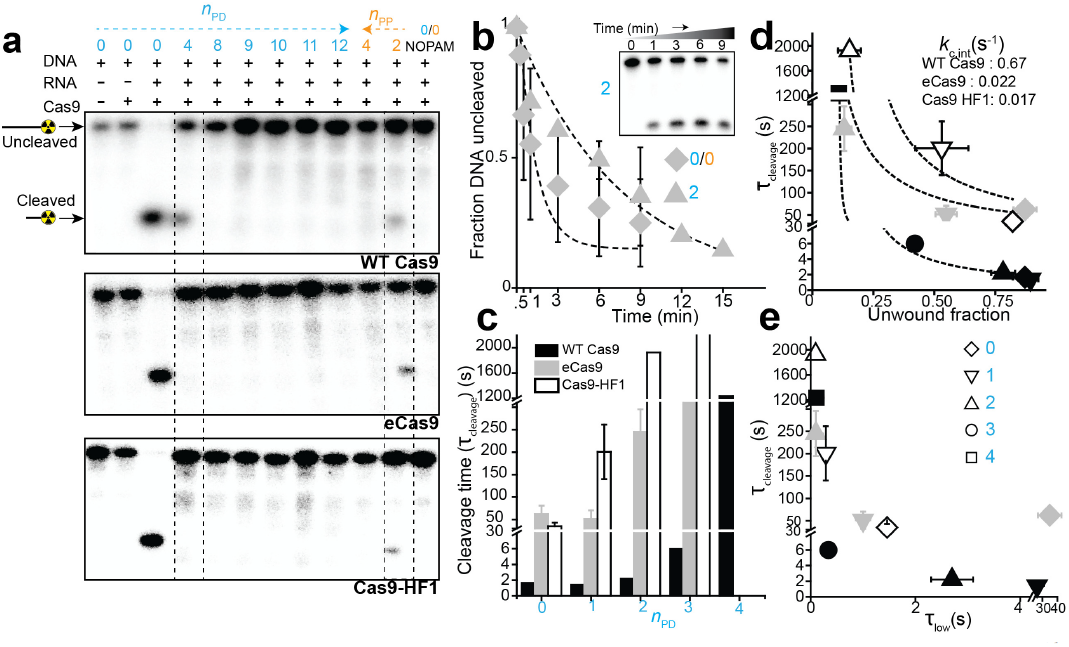
Cleavage vs mismatches and its relation with DNA unwinding. **(a)** Cas9-RNA induced cleavage pattern analyzed by 15% denaturing polyacrylamide gel electrophoresis of target strand 5_’_- labeled with ^32-^P. Dashed boxes highlight DNA targets with *n*_PD_ =4 and *n*_PP_ =2 that show differences between EngCas9s and WT Cas9. **(b)** Fraction of uncleaved DNA vs time for eCas9 as a function of time and single exponential decay fits to obtain cleavage time *τ*_cleavage_. Inset shows a representative gel image. [DNA] = 1 nM. [Cas9-RNA] = 100 nM. **(c)** *τ*_cleavage_ vs *n*_PD_. Data for WT Cas9-RNA is taken from a previous study^28^. **(d)** *τ*_cleavage_ vs unwound fraction as shown in Fig. 2c. Fits are made to determine *k*_c,int_ (see text). (e) *τ*_cleavage_ vs. unwound state lifetime as shown in **Fig 2d**. *n*_PD_ and *n*_PP_ are shown in cyan and orange, respectively.

HNH and RuvC nuclease domains cleave target and non-target strand respectively. Additional 3_’_-5_’_ exonuclease activity of RuvC^3,25,27^, which normally causes multiple bands of cleaved non-target strand, decreased with increasing *n*_PD_, and more efficiently so for EngCas9s (**Supplementary Fig. 8**).

Armed with the kinetic information on DNA binding, interrogation, unwinding/rewinding and cleavage, we can clarify in which reaction steps sequence mismatches and Cas9 mutations exert their influence. Because cleavage likely occurs from the unwound state, we estimated the lower limits of intrinsic rate of cleavage (*k*_c,int_) from the apparent cleavage time *τ*_cleavage_ and unwound fraction by fitting *τ*_cleavage_ vs unwound fraction using *τ*_cleavage_ = 1/([unwound fraction]\**k*_c,int_+C). Because a single value of *k*_c,int_ was able to fit the data for each Cas9, we conclude that changes in *τ*_cleavage_ caused by mismatches can be attributed to changes in the unwound fraction (**Fig. 3d**). Therefore, although DNA molecules with different *n*_PD_ values show different extents of unwinding, their unwound states have similar intrinsic cleavage rates. *k*_c,int_ was ~0.67 s^-1^ for WT Cas9 but decreased to 0.025-0.015 s^-1^ for EngCas9s.

Inserting two single bp mismatches in the PAM-distal region gave a predominantly rewound state with only brief excursions to the unwound state for all Cas9s (**Fig. 4a**), indicating that there is no difference in unwound fraction among them. Nevertheless, WT Cas9 cleaved the DNA whereas EngCas9s did not, further confirming the much higher *k*_c,int_ for WT Cas9 which in turn allows cleavage even from a transiently populated unwound state, likely contributing to higher off-target cleavage yields (**Fig. 4b**).

**Figure 4.**
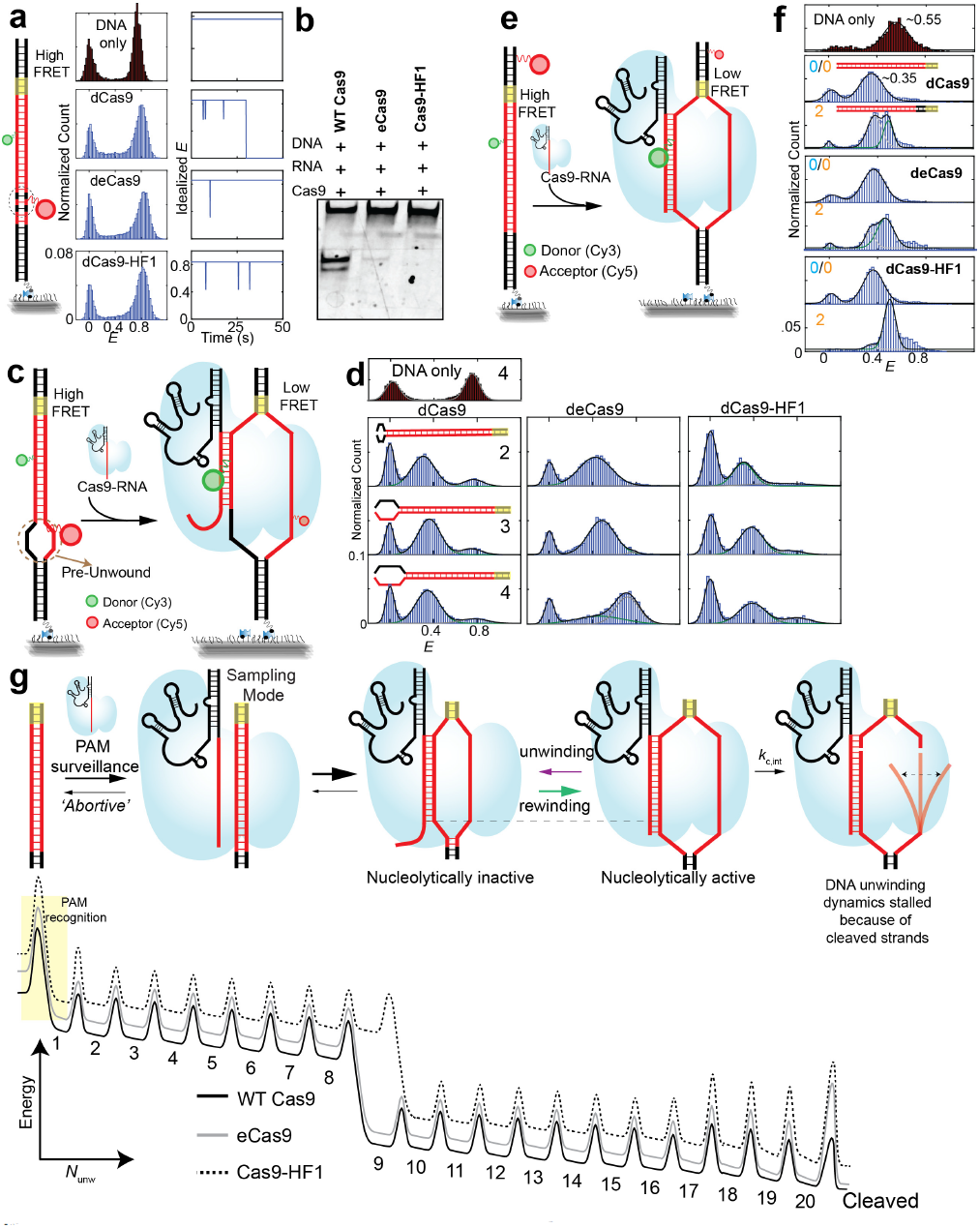
Cas9-RNA induced unwinding of various DNA and mechanisms of increased specificity by EngCas9s. **(a)** Unwinding and cleavage of DNA with internal single base mismatches at positions 16^th^ and 18^th^. *E* histograms were obtained at [Cas9-RNA] =100 nM or in its absence. Idealized *E* traces show transient unwinding observed in a subset of molecules. **(b)** Gel image of non-target strand shows that WT Cas9, but not EngCas9s, cleave the DNA. **(c)** Schematics of Cas9-RNA induced unwinding of pre-unwound DNA. **(d)** *E* histograms obtained at [Cas9-RNA] =100 nM or in its absence. Black numbers denote the number of base pairs pre-unwound and are mismatched. **(e)** Schematics of Cas9-RNA induced unwinding of PAM-proximal region. **(f)** *E* histograms obtained at [Cas9-RNA] =100 nM or in its absence. *n*_PP_ is shown in orange digits. **(g)** Proposed model of DNA targeting, unwinding, rewinding, and cleavage (top). Energy diagram as a function of *N*_unw_, the number of unwound base pairs.

### Mechanism of mismatch sensitivity of EngCas9s

The changing balance between unwinding and rewinding caused by mismatches reflect the energetic competition between target strand annealing with RNA vs with non-target strand. Mismatches would favor rewinding to recover the parental duplex by disrupting RNA/DNA duplex. We found that mutations in EngCas9s further promote rewinding, helping improve cleavage specificity. In order to gain additional insights into how EngCas9s more readily rewind in the presence of mismatches, we disrupted the parental DNA duplex by pre-unwinding the PAM-distal mismatched portion (**Fig. 4c**). Pre-unwinding shifted the balance toward unwinding for WT Cas9 and Ca9-HF1 (**Fig. 4d**), possibly because the residues that sequester the non-target strand, mutated in eCas9, are still intact. The result also explains why pre-unwinding allows rapid cleavage of mismatched DNA^28^. In contrast, eCas9 failed to shift *E* distribution toward a low value in the presence of PAM-distal mismatches (**Fig. 4d & Supplementary Fig. 16**), likely because the residues that help sequester the unwound non-template strand are mutated (**Fig. 1b & Supplementary Fig. 17**).

### PAM-proximal DNA unwinding

To investigate if EngCas9-RNA mutations also affect PAM-proximal DNA unwinding we designed a DNA construct with a donor and an acceptor at the PAM proximal site separated by 12 bp (**Fig. 4e & Supplementary Fig. 18**). Cas9-RNA binding to cognate DNA shifted *E* from 0.55 to 0.35 for all Cas9s, indicating a stable DNA unwinding at PAM-proximal site. With *n*_PP_ = 2, the fraction of unwound population decreased to 50% for dCas9, 40% for deCas9 and < 5% for dCas9-HF1 (**Fig. 4f**), mirroring the decreases in *f*_bound_. However, this decrease did not correlate with cleavage efficiency of DNA target with *n*_PP_ = 2, where EngCas9 were more efficient than WT Cas9 (**Fig. 3a**). Therefore, EngCas9s can cleave DNA better once the initial barrier caused by PAM-proximal mismatches is overcome.

## Discussion

Like WT Cas9, EngCas9s also require only 9-10 PAM-proximal matches for ultra-stable binding and thus will likely not confer significant specificity advances for applications utilizing Cas9 binding, for example transcription regulation^29^ or imaging^30^. EngCas9s, however, do show a modest increase in binding specificity. For example, Cas9-HF1 requires one additional basepair match for ultra-stable binding compared to WT Cas9. Ultra-stable binding to partially matching sequences can sequester Cas9-RNA, limiting the speed of genome editing. To overcome sequestration, one would need higher Cas9-RNA concentrations which may in turn lead to an increase in off-target cleavage. One base pair difference would mean about four-fold reduction in such off-target sinks for Cas9-HF1, and even such a modest improvement in binding specificity may improve cleavage specificity by reducing the total concentration of Cas9-RNA required.

Cas9_’_s cleavage activity requires many more sequence matches than stable binding^3,4^ and we found that DNA unwinding, instead of DNA binding, is strongly correlated with the cleavage rate when we vary the DNA sequence or Cas9 mutations. Unwound DNA configuration is likely verified by REC3 domain and linker connecting HNH and RuvC nuclease domains in Cas9^21,31^, guiding HNH nuclease movement for cleavage. REC2 (coupled to REC3) proofreading and HNH movements are impaired by PAM-distal mismatches^18,21^ that we show here to also impair DNA unwinding. DNA unwinding occurs even in the absence of divalent cations (**Supplementary Fig. 19**) whereas HNH movement requires divalent cations^18^. Therefore, DNA unwinding is likely an upstream trigger of HNH movement and does not occur with or because of HNH movement as had been suggested in a simulation study^31^. This also means that DNA unwinding is the critical step that verifies sequence matching, which then subsequently triggers a series of steps toward cleavage including HNH movement.

The impact of PAM-distal matches in opposing DNA unwinding is even greater for EngCas9 because their mutations destabilize the unwound DNA configuration, and because cleavage likely proceeds from the unwound state, the increased sensitivity to mismatches in DNA unwinding is likely to contribute to the increased specificity of cleavage by EngCas9s. Any perturbation that destabilizes unwound DNA or fully extended R-loop will delay or fully impair the cleavage action and is also likely the reason why Cas9 with truncated guide-RNAs are more specific in cleavage^32^.

DNA targets with only a few interspersed single base-pair mismatches at PAM-distal site can prevent stably unwound states by all Cas9 but even transiently unwound states appear to allow cleavage by WT Cas9, but not by EngCas9s which we showed here to have much lower intrinsic cleavage rate. Therefore, the large enhancement in cleavage specificity for EngCas9s may in part be due to the lower intrinsic cleavage rate that prevents cleavage from transiently unwound states that are populated in partially matching sequences.

**Fig. 4g** summarizes our findings on the three possible mechanisms of specificity increase for EngCas9 in the form of free energy diagram along the axis of *N*_unw_, the number of bp unwound, leading toward cleavage. Once the initial barrier of PAM detection is overcome, DNA unwinding proceeds in the direction from PAM-proximal to PAM-distal. There is a sudden drop in energy at *N*_unw_=9 for WT Cas9 and eCas9, marking a threshold for stable binding. Because the threshold is shifted to *N*_unw_=10 for Cas9-HF1, Cas9-HF1 can achieve higher cleavage specificity by reducing the number of off-target sites which can otherwise sequester Cas9-RNA (Mechanism 1). EngCas9s also tilt the energetic balance toward rewinding in the PAM-distal region such that PAM-distal mismatches readily depopulate the fully unwound state needed for cleavage (Mechanism 2). Finally, EngCas9s have lower intrinsic cleavage rates from the unwound state, represented as higher energetic barrier against cleavage, preventing cleavage from transiently unwound off-targets (Mechanism 3).

EngCas9s still cleave DNA targets with ~3 PAM-distal mismatches thus there is room for further improvement. Cas9-HF1 and eCas9 mutations eliminated some of the favorable sequence-independent interactions for target and non-target strands separately and may act in an independent manner and thus, their mutations may be combined. EngCas9 mutations may also be combined with truncated RNA to improve the specificity even further. But, combining a number of such strategies that destabilize unwound state and decrease the intrinsic cleavage rate from the unwound state as we showed here could lead to substantially reduced on-target cleavage as well, as has been observed when Cas9-HF1 was used with the truncated RNA^10^.

## Materials and methods

### Preparation of DNA targets

DNA targets used in smFRET assay for DNA interrogation by Cas9-RNA are the same as what were used previously for WT Cas9 studies ^9^ and **Supplementary Figure 1** shows the overall schematics. The schematic of DNA targets used for smFRET assay of Cas9-RNA induced DNA unwinding is shown in **Supplementary Figure 7, 12, 18**. All DNA oligonucleotides were purchased from Integrated DNA Technologies (IDT, Coralville, IA 52241). A thymine modified with an amine group through a C6 linker (amino-dT) was used to label DNA with Cy3 or Cy5 N-hydroxysuccinimido (NHS). Non-target strand, target strand and a 22 nt biotinylated adaptor strand were assembled by mixing them in buffered solution with 10 mM Tris-HCl, pH 8 and 50 mM NaCl and heating to 90 °C followed by cooling to room temperature over 3 hrs. For DNA unwinding by surface-tethered Cas9-RNA, the biotin adaptor strand was omitted. Full sequence and modifications of DNA targets used in smFRET assays for DNA interrogation and DNA unwinding are shown in **Supplementary Table 1** and **Supplementary Table 2**, respectively. For radio-labeled gel electrophoresis cleavage experiments, the target strand was phosphorylated with P^32^ using T4 polynucleotide kinase reaction and was annealed with the non-target strand as described above. Sequences are the same as in **Supplementary Table 1** and **Supplementary Table 2** but without the biotinylated adaptor strand. For fluorescently-labeled gel electrophoresis cleavage experiments, the DNA targets used are the same as those used in smFRET assays.

### Expression and purification of Cas9

All Cas9 were expressed and purified as described previously^2,33^. A pET-based expression vector was used for protein expression which consisted of sequence encoding Cas9 (Cas9 residues 1-1368 from *S. pyogenes*) and an N-terminal decahistidine-maltose binding protein (His10-MBP) tag, followed by a peptide sequence containing a tobacco etch virus (TEV) protease cleavage site.

Mutations for cleavage impairment, enhanced mutations (eCas9), HF1 mutations or the desired combinations of them were introduced by site-directed mutagenesis (QuickChange Lightning; Agilent Technologies, Santa Clara, CA 95050). Proteins were expressed in *E. coli* strain BL21 Rosetta 2 (DE3) (EMD Biosciences), grown in TB (Terrific Broth) or 2YT medium (higher expression obtained for TB) at 37 °C for a few hours. When the optical density at 600 nm (OD_600_) reached 0.6, protein expression was induced with 0.5 mM IPTG and the temperature was lowered to 18 °C. The induction was then continued for 12-16 h. The medium was then discarded and cells were harvested. The harvested cells were lysed in 50 mM Tris pH 7.5, 500 mM NaCl, 5% glycerol, 1 mM TCEP, supplemented with protease inhibitor cocktail (Roche) and with/without Lysozyme (Sigma Aldrich), and then homogenized in a microfluidizer (Avestin) or homogenized with Fisher Model 500 Sonic Dismembrator (Thermo Fisher Scientific) at 30 % amplitude in 3 one minute cycles, each consisting of series of 2 s sonicate-2 s repetitions. The lysed solution was then ultra-centrifuged at 15,000 *g* for 30-45 minutes, supernatant of lysate was collected and cellular debris was discarded. The supernatant was added to Ni-NTA agarose resin (Qiagen). The resin was washed extensively with 50 mM Tris pH 7.5, 500 mM NaCl, 10 mM imidazole, 5% glycerol, 1 mM TCEP and the bound protein was eluted in a single-step with 50 mM Tris pH 7.5, 500 mM NaCl, 300 mM imidazole, 5% glycerol, 1 mM TCEP. Dialysis of Cas9 into Buffer A (20 mM TrisCl pH 7.5, 125 mM KCl, 5% glycerol, 1 mM TCEP) and cleavage of TEV-protease site by TEV protease was simultaneously carried out overnight at 4 °C. Deconstitution of 10-His-MBP-TEV-Cas9 by TEV protease resulted in 10-His-MBP and Cas9 constituents in the solution. Another round of NiNTA agarose column was performed to arrest 10-His-MBP out of the solution and obtain free Cas9. Cas9 was then further purified by size-exclusion chromatography on a Superdex 200 16/60 column (GE Healthcare) in Cas9 Storage Buffer (20 mM Tris-Cl pH 7.5, 100 mM KCl, 5% glycerol and 5 mM MgCl_2_) and stored at -80 °C. All the purification steps were performed at 4 °C. In some preparations, TEV protease was first added to the elutant and cleavage of the protein fusion was carried out overnight. Following TEV protease cleavage, Cas9 was then dialyzed into Buffer A (20 mM Tris-Cl pH 7.5, 125 mM KCl, 5% glycerol, 1 mM TCEP) for 3 h at 4 °C, before being applied onto a 5 ml HiTrap SP HP sepharose column (GE Healthcare). After washing with Buffer A for three column volumes, Cas9 was eluted using a linear gradient from 0-100% Buffer B (20 mM Tris-Cl pH 7.5, 1 M KCl, 5% glycerol, 1 mM TCEP) over 20 column volumes. The protein was further purified by gel filtration chromatography on a Superdex 200 16/60 column (GE Healthcare) in Cas9 Storage Buffer (20 mM Tris-Cl pH 7.5, 200 mM KCl, 5% glycerol, 1 mM TCEP). Cas9 was stored at -80 °C.

### Preparation of guide-RNA and Cas9-RNA

Guide-RNA for Cas9 is a combination of CRISPR RNA (crRNA) and trans-activating crRNA (tracrRNA). For smFRET assay for DNA interrogation by Cas9-RNA, crRNA with an amino-dT was purchased from IDT and labeled with Cy5-NHS. Location of the Cy5 in the crRNA is shown in **Supplementary Figure 1.** The tracrRNA was *in vitro* transcribed as described previously^4^. For DNA unwinding smFRET assay, both crRNA and tracrRNA were unlabeled and transcribed *in vitro.* Guide-RNA was assembled by mixing crRNA and tracrRNA at 1:1.2 ratio in buffer containing 10 mM Tris HCl (pH 8) and 50 mM NaCl, heated to 90 °C and slowly cooled to room temperature. Cas9-RNA was assembled by mixing guide-RNA and Cas9 at a ratio of 1:3 in Cas9-RNA activity buffer (20 mM Tris HCl (pH 8), 100 mM KCl, 5 mM MgCl_2_, 5% v/v glycerol). Cas9-RNA cleavage activity on cognate sequence used for smFRET assay for DNA interrogation was characterized previously^4^. We used a slightly difference sequence for optical placement of amino-dTs for DNA unwinding smFRET assay and Cas9-RNA cleavage activity on that sequence was also confirmed (**Supplementary Fig. 8**). RNA sequences are available in **Supplementary Table 1 & 2.** For smFRET assay of DNA unwinding by surface-tethered Cas9-RNA, a biotin adaptor DNA strand was annealed to a complementary extension on guide-RNA (**Supplementary Figure 12**)

### Single-molecule fluorescence imaging and data analysis

DNA targets were immobilized on the polyethylene glycol-passivated flow chamber surface (purchased from Johns Hopkins University Microscope Supplies Core or prepared following protocols reported previously ^22^ using neutravidin-biotin interaction and imaged in the presence of Cas9-RNA at the stated concentration using two-color total internal reflection fluorescence microscopy. For DNA unwinding by surface-tethered Cas9-RNA smFRET assay, 20 nM of biotin-labeled Cas9-RNA was immobilized on surface before adding FRET pair labeled DNA targets. All the imaging experiments were done at room temperature in a Cas9-RNA activity buffer with oxygen scavenging for extending photostability (20 mM Tris-HCl, 100 mM KCl, 5 mM MgCl_2_, 5% (v/v) glycerol, 0.2 mg ml^-1^ BSA, 1 mg ml^-1^ glucose oxidase, 0.04 mg ml^-1^ catalase, 0.8% dextrose and saturated Trolox (>5 mM)^34^. Time resolution was 100 ms unless stated otherwise. Detailed methods of single-molecule FRET (smFRET) data acquisition and analysis have been described previously^22^. Video recordings obtained using EMCCD camera (Andor) were processed to extract single molecule fluorescence intensities at each frame and custom written scripts were used to calculate FRET efficiencies. Data acquisition and analysis software can be downloaded from https://cplc.illinois.edu/software/. FRET efficiency (*E*) of the detected spot was approximated as *E* = IA/(*I*_D_+*I*_A_), where *I*_D_ and *I*_A_ are background and leakage corrected emission intensities of the donor and acceptor, respectively.

### *E* histograms, Cas9-RNA bound DNA fraction and unwound fraction

Unless stated otherwise, first five frames of each molecule_’_s *E* value time traces were used as data points to construct *E* histograms. At least 2,500 molecules were included in each histogram. Cas9-RNA bound DNA fraction was calculated as a fraction of data points with *E* > 0.75 and was then normalized relative to Cas9-RNA bound DNA fraction for the cognate DNA target, which is not 100% because missing or inactive acceptor fluorophore, and is referred to as *f*_bound_. To estimate *K*_*d*_, *f*_bound_ vs Cas9-RNA concentration (*c*) was fit using *f*_bound_ = M × c / (*K*_*d*_ + c) where M is the maximum observable *f*_bound_. M is typically less than 1 because inactive or missing acceptors or because not all of the DNA on the surface are capable of binding Cas9-RNA. For DNA unwinding smFRET assay, unwound fraction was calculated as the ratio of data points with 0.2 <*E* <0.6 and total number of data points with *E>0.2.*

### Kinetic analysis of DNA interrogation by Cas9-RNA

A bimolecular association/dissociation kinetics was used for the analysis of DNA binding by Cas9-RNA. DNA Target + Cas9-RNA [inline1] Cas9-RNA-DNA

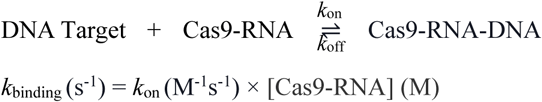

A hidden Markov model analysis of smFRET traces of DNA targets that showed real-time reversible association/dissociation of Cas9-RNA yielded three pre-dominant FRET states, of zero, mid and high *E* values. Dwell times of high FRET (*E*>0.6) states before their transition to zero FRET (*E*<0.2) states were used to calculate lifetime of high FRET (*τ*_high_) (**Supplementary Fig. 5c**) binding events by fitting their distribution using a double-exponential decay. Dwell times of mid FRET (0.2 <*E* <0.6) states before their transition to zero FRET (*E*<0.2) states were used to calculate lifetime of mid FRET (*τ*_mid_) (**Supplementary Fig. 5d**) binding events by fitting their distribution using a single exponential decay.

To calculate the total bound state lifetime (*τ*_avg_) (**Fig. 1d**), mid and high FRET states were taken as bound states and zero FRET state as the unbound state. The survival probability of all the bound state events (mid and high FRET states taken as a single state, FRET > 0.2) vs time was fit using a double exponential decay profile (*A*_1_ *exp*(-*t*/*τ*_1_) +*A*_2_ *exp*(-*t*/*τ*_2_)). *τ*_avg_ is an amplitude weighted average of two distinct lifetimes *τ*_1_ and *τ*_2_.

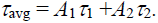

*A*_1_ values are also used as the fraction of mid FRET state binding events (*f*_mid FRET_). Dwell time distribution of zero FRET state (*E*<0.2) was fit to a single exponential decay to calculate lifetime of unbound state (*τ*_unbound_) (**Fig. 1f**). Inverse of *τ*_unbound_ was used to calculate *k*_binding_ and *k*_on_. Since Cas9-RNA DNA association were sequence independent, a mean of *k*_on_ for different DNA target was used to determine *k*_on_ for each Cas9 (**Fig. 1f**)

### Kinetic analysis of Cas9-RNA induced DNA unwinding and rewinding

Cas9-RNA induced DNA unwinding was modeled as a two-state system (**Fig. 2b**):

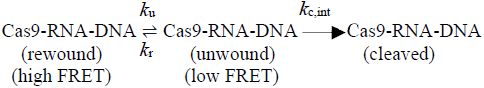

A hidden Markov model analysis segmented single molecule time traces into unwound and rewound states (low and high FRET states). Dwell time histograms were fitted using single exponential decay to determine the average lifetime of the high (*τ*_high_) and low FRET (*τ*_low_) states respectively (**Fig. 2d-e**). A small fraction of its smFRET time trajectories had a constant high FRET without any fluctuations, likely representing DNA molecules unable to bind Cas9-RNA and were excluded from kinetic analysis. Unwinding rate (*k*_u_) was calculated as inverse of *τ*_high_ and rewinding rate (*k*_r_) was calculated as inverse of *τ*_low._ Intrinsic cleavage rate *k*_c,int_ was determined by fitting cleavage lifetime vs unwound fraction using *τ*_cleavage_ = 1/([unwound fraction]\**k*_c,int_ + C), where C=0 for eCas9 but an extremely small value of C had to be used for WT Cas9 and Cas9-HF1.

In smFRET experiments to capture the initial DNA unwinding event, distribution of dwell times of the initial high FRET state before the first transition to the low FRET state was fit with a single exponential to obtain *τ*_unwinding_.

### Gel-electrophoresis to investigate Cas9-RNA induced cleavage

FRET pair labeled DNA targets used in the smFRET DNA unwinding assays were also used in fluorescence-based gel electrophoresis experiments. The DNA targets were incubated with Cas9-RNA in Cas9-RNA activity buffer (20 mM Tris HCl (pH 8), 100 mM KCl, 5 mM MgCl_2_, 5% v/v glycerol) at the specified concentrations and for specified durations. The reaction samples were then denatured by formamide loading buffer (95% formamide, 5 mM EDTA) and resolved via polyacrylamide gel electrophoresis (PAGE) using 15% Polyacrylamide TBE Urea pre-cast gels (Bio-rad laboratories) and imaged via Cy5/Cy3 excitation (GE Amersham Imager 600). Sequences of DNA targets and guide-RNA are shown in **Supplementary Table 2**.

For radio-labeled PAGE experiments, P^32^ -labeled DNA targets at 1 nM concentration were incubated with 100 nM Cas9-RNA in Cas9-RNA activity buffer (20 mM Tris HCl (pH 8), 100 mM KCl, 5 mM MgCl_2_, 5% v/v glycerol) at for 10 minutes. The reaction samples were then denatured by formamide loading buffer (95% formamide, 5 mM EDTA), resolved via PAGE and imaged via phosphorimaging (GE Healthcare). For the rate of cleavage experiments, aliquots of Cas9-RNA DNA running reaction were taken at different time points for the PAGE analysis. Sequences of DNA targets and guide-RNA are the same as in **Supplementary Table 2** except that the biotinylated adaptor strand is omitted. *τ*_cleavage_ for WT Cas9 is taken from a study^28^ which used a slightly different protospacer, shown in **Supplementary Table 1.** In a control, the *τ*_cleavage_ for WT Cas9 were similar between the two protospacers and much lower than *τ*_cleavage_ for EngCas9.

## Acknowledgements

The project was supported by grants from National Science Foundation (PHY-1430124 to T.H.) and National Institutes of Health (GM065367; GM112659 to T.H and GM097330 to S.B.); T.H. is an investigator with the Howard Hughes Medical Institute. J.M is supported by the Nation Institutes of Health Chemical Biology Interface training program (T32GM080189). We would like to thank Jennifer A. Doudna and Samuel H. Sternberg for useful early discussions about the design of experiments. We would also like to thank Samuel H. Sternberg and Janice S. Chen of Doudna lab for generously providing some Cas9 stocks and EngCas9 expression plasmids, respectively.

## Author contributions and Competing interests

D.S. and T.H. designed the experiments. D.S. and Y.W. performed smFRET DNA interrogation experiments. D.S. performed smFRET DNA unwinding experiments. D.S. and J.M. performed gel-based experiments. D.S., J.M. expressed and purified Cas9s. D.S., Y.W., O.Y., J.F., A.P., and D.C. performed or helped with the data analysis. O.Y. assisted with some experiments. A.P. assisted with PEG passivation of some slides. D.S., T.H., and S.B. discussed the data. D.S. and T.H. wrote the manuscript.

Authors declare no competing interests.

## References

1 Marraffini, L. A. & Sontheimer, E. J. CRISPR interference: RNA-directed adaptive immunity in bacteria and archaea. Nature reviews. Genetics 11, 181–190, doi:10.1038/nrg2749 (2010).

2 Gasiunas, G., Barrangou, R., Horvath, P. & Siksnys, V. Cas9–crRNA ribonucleoprotein complex mediates specific DNA cleavage for adaptive immunity in bacteria. Proceedings of the National Academy of Sciences of the United States of America 109, E2579–E2586, doi:10.1073/pnas.1208507109 (2012).

3 Jinek, M., Chylinski, K., Fonfara, I., Hauer, M., Doudna, J. A. & Charpentier, E. A programmable dual-RNA-guided DNA endonuclease in adaptive bacterial immunity. Science 337, 816–821, doi:10.1126/science.1225829 (2012).

4 Sternberg, S. H., Redding, S., Jinek, M., Greene, E. C. & Doudna, J. A. DNA interrogation by the CRISPR RNA-guided endonuclease Cas9. Nature 507, 62–67, doi:10.1038/nature13011 (2014).

5 Wang, H., La Russa, M. & Qi, L. S. CRISPR/Cas9 in Genome Editing and Beyond. Annual review of biochemistry 85, 227–264, doi:10.1146/annurev-biochem-060815-014607 (2016).

6 Barrangou, R. & Doudna, J. A. Applications of CRISPR technologies in research and beyond. Nature biotechnology 34, 933–941, doi:10.1038/nbt.3659 (2016).

7 Wu, X., Kriz, A. J. & Sharp, P. A. Target specificity of the CRISPR-Cas9 system. Quantitative biology 2, 59–70, doi:10.1007/s40484-014-0030-x (2014).

8 O’Geen, H., Yu, A. S. & Segal, D. J. How specific is CRISPR/Cas9 really? Current opinion in chemical biology 29, 72–78, doi:10.1016/j.cbpa.2015.10.001 (2015).

9 Singh, D., Sternberg, S. H., Fei, J., Doudna, J. A. & Ha, T. Real-time observation of DNA recognition and rejection by the RNA-guided endonuclease Cas9. Nature communications 7, 12778, doi:10.1038/ncomms12778 (2016).

10 Kleinstiver, B. P., Pattanayak, V., Prew, M. S., Tsai, S. Q., Nguyen, N. T., Zheng, Z. & Joung, J. K. High-fidelity CRISPR-Cas9 nucleases with no detectable genome-wide off-target effects. Nature 529, 490–495, doi:10.1038/nature16526 (2016).

11 Slaymaker, I. M., Gao, L., Zetsche, B., Scott, D. A., Yan, W. X. & Zhang, F. Rationally engineered Cas9 nucleases with improved specificity. Science 351, 84–88, doi:10.1126/science.aad5227 (2016).

12 Joo, C., Balci, H., Ishitsuka, Y., Buranachai, C. & Ha, T. Advances in single-molecule fluorescence methods for molecular biology. Annual review of biochemistry 77, 51–76, doi:10.1146/annurev.biochem.77.070606.10 1543 (2008).

13 Sternberg, S. H., Redding, S., Jinek, M., Greene, E. C. & Doudna, J. A. DNA interrogation by the CRISPR RNA-guided endonuclease Cas9. Nature 507, 62–67, doi:10.1038/nature13011 (2014).

14 Szczelkun, M. D., Tikhomirova, M. S., Sinkunas, T., Gasiunas, G., Karvelis, T., Pschera, P., Siksnys, V. & Seidel, R. Direct observation of R-loop formation by single RNA-guided Cas9 and Cascade effector complexes. Proceedings of the National Academy of Sciences of the United States of America 111, 9798–9803, doi:10.1073/pnas.1402597111 (2014).

15 Rutkauskas, M., Sinkunas, T., Songailiene, I., Tikhomirova, M. S., Siksnys, V. & Seidel, R. Directional R-Loop Formation by the CRISPR-Cas Surveillance Complex Cascade Provides Efficient Off-Target Site Rejection. Cell reports, doi:10.1016/j.celrep.2015.01.067 (2015).

16 Redding, S., Sternberg, S. H., Marshall, M., Gibb, B., Bhat, P., Guegler, C. K., Wiedenheft, B., Doudna, J. A. & Greene, E. C. Surveillance and processing of foreign DNA by the Escherichia coli CRISPR-Cas system. Cell 163, 854–865, doi:10.1016/j.cell.2015.10.003 (2015).

17 Blosser, T. R. Two distinct DNA binding modes guide dual roles Of a CRISPR-Cas protein complex. 58, 60–70, doi:10.1016/j.molcel.2015.01.028 (2015).

18 Dagdas, Y. S., Chen, J. S., Sternberg, S. H., Doudna, J. A. & Yildiz, A. A conformational checkpoint between DNA binding and cleavage by CRISPR-Cas9. Science Advances 3, doi:10.1126/sciadv.aao0027 (2017).

19 Josephs, E. A., Kocak, D. D., Fitzgibbon, C. J., McMenemy, J., Gersbach, C. A. & Marszalek, P. E. Structure and specificity of the RNA-guided endonuclease Cas9 during DNA interrogation, target binding and cleavage. Nucleic acids research 43, 8924–8941, doi:10.1093/nar/gkv892 (2015).

20 Lim, Y., Bak, S. Y., Sung, K., Jeong, E., Lee, S. H., Kim, J. S., Bae, S. & Kim, S. K. Structural roles of guide RNAs in the nuclease activity of Cas9 endonuclease. Nature communications 7, 13350, doi:10.1038/ncomms13350 (2016).

21 Chen, J. S., Dagdas, Y. S., Kleinstiver, B. P., Welch, M. M., Sousa, A. A., Harrington, L. B., Sternberg, S. H., Joung, J. K., Yildiz, A. & Doudna, J. A. Enhanced proofreading governs CRISPR-Cas9 targeting accuracy. Nature, doi:10.1038/nature24268 (2017).

22 Roy, R., Hohng, S. & Ha, T. A practical guide to single-molecule FRET. Nature methods 5, 507–516, doi:10.1038/nmeth.1208 (2008).

23 McKinney, S. A., Joo, C. & Ha, T. Analysis of Single-Molecule FRET Trajectories Using Hidden Markov Modeling. Biophysical Journal 91, 1941–1951, doi:10.1529/biophysj.106.082487 (2006).

24 Boyle, E. A., Andreasson, J. O. L., Chircus, L. M., Sternberg, S. H., Wu, M. J., Guegler, C. K., Doudna, J. A. & Greenleaf, W. J. High-throughput biochemical profiling reveals sequence determinants of dCas9 off-target binding and unbinding. Proceedings of the National Academy of Sciences of the United States of America 114, 5461–5466, doi:10.1073/pnas.1700557114 (2017).

25 Jiang, F., Taylor, D. W., Chen, J. S., Kornfeld, J. E., Zhou, K., Thompson, A. J., Nogales, E. & Doudna, J. A. Structures of a CRISPR-Cas9 R-loop complex primed for DNA cleavage. Science 351, 867–871, doi:10.1126/science.aad8282 (2016).

26 Richardson, C. D., Ray, G. J., DeWitt, M. A., Curie, G. L. & Corn, J. E. Enhancing homology-directed genome editing by catalytically active and inactive CRISPRCas9 using asymmetric donor DNA. Nature biotechnology 34, 339–344, doi:10.1038/nbt.3481 (2016).

27 Zuo, Z. & Liu, J. Cas9-catalyzed DNA Cleavage Generates Staggered Ends: Evidence from Molecular Dynamics Simulations. Scientific reports 5, 37584, doi:10.1038/srep37584 (2016).

28 Sternberg, S. H., LaFrance, B., Kaplan, M. & Doudna, J. A. Conformational control of DNA target cleavage by CRISPR-Cas9. Nature 527, 110–113, doi:10.1038/nature15544 (2015).

29 Gilbert, L. A., Larson, M. H., Morsut, L., Liu, Z., Brar, G. A., Torres, S. E., Stern-Ginossar, N., Brandman, O., Whitehead, E. H., Doudna, J. A., Lim, W. A., Weissman, J. S. & Qi, L. S. CRISPR-Mediated Modular RNA-Guided Regulation of Transcription in Eukaryotes. Cell 154, 442–451, doi:10.1016/j.cell.2013.06.044 (2013).

30 Chen, B., Gilbert, L. A., Cimini, B. A., Schnitzbauer, J., Zhang, W., Li, G. W., Park, J., Blackburn, E. H., Weissman, J. S., Qi, L. S. & Huang, B. Dynamic Imaging of Genomic Loci in Living Human Cells by an Optimized CRISPR/Cas System. Cell 155, 1479–1491, doi:10.1016/j.cell.2013.12.001 (2013).

31 Palermo, G., Miao, Y., Walker, R. C., Jinek, M. & McCammon, J. A. CRISPR-Cas9 conformational activation as elucidated from enhanced molecular simulations. Proceedings of the National Academy of Sciences of the United States of America 114, 7260–7265, doi:10.1073/pnas.1707645114 (2017).

32 Fu, Y., Sander, J. D., Reyon, D., Cascio, V. M. & Joung, J. K. Improving CRISPR-Cas nuclease specificity using truncated guide RNAs. Nature biotechnology 32, 279–284, doi:10.1038/nbt.2808 (2014).

33 Jinek, M., Jiang, F., Taylor, D. W., Sternberg, S. H., Kaya, E., Ma, E., Anders, C., Hauer, M., Zhou, K., Lin, S., Kaplan, M., Iavarone, A. T., Charpentier, E., Nogales, E. & Doudna, J. A. Structures of Cas9 endonucleases reveal RNA-mediated conformational activation. Science 343, 1247997, doi:10.1126/science.1247997 (2014).

34 Rasnik, I., McKinney, S. A. & Ha, T. Nonblinking and long-lasting single-molecule fluorescence imaging. Nature methods 3, 891–893, doi:10.1038/nmeth934 (2006).

